# Structural robustness of mammalian transcription factor networks reveals plasticity across development

**DOI:** 10.1101/209528

**Authors:** JL Caldu-Primo, ER Alvarez-Buylla, J Davila-Velderrain

## Abstract

Network biology aims to understand cell behavior through the analysis of underlying complex biomolecular networks. Inference of condition-specific interaction networks from epigenomic data enables the characterization of the structural plasticity that regulatory networks can acquire in different tissues of the same organism. From this perspective, uncovering specific patterns of variation by comparing network structure among tissues could provide insights into systems-level mechanisms underlying cell behavior. Following this idea, here we propose an empirical framework to analyze mammalian tissue-specific networks, focusing on characterizing and contrasting their structure and behavior in response to perturbations. We structurally represent the state of the cell/tissue by condition specific transcription factor networks generated using chromatin accessibility data, and we profile their systems behavior in terms of the structural robustness against random and directed perturbations. Using this framework, we unveil the structural heterogeneity existing among tissues at different levels of differentiation. We uncover a novel and conserved systems property of regulatory networks underlying embryonic stem cells (ESCs): in contrast to terminally differentiated tissues, the promiscuous regulatory connectivity of ESCs produces a globally homogeneous network resulting in increased structural robustness. Possible biological consequences of this property are discussed.

## Introduction

A central tenet of systems biology is that cell behavior can be understood in terms of the structure and dynamics of underlying complex molecular networks.^1, 2^ Under such paradigm, major efforts have been made to systematically map and characterize the properties of molecular networks at different levels of organization. Reference protein-protein interaction, metabolic, and transcriptional regulatory networks have been constructed and are being frequently updated in several model organisms.^3–5^ Initial efforts have largely focused on providing an organismal reference for the global network structure.

Network theory provides methods for the systemic description of a network’s structure and its dynamics.^6–8^ One of the major results of network biology is the discovery within the reference networks of apparently universal organizational properties across the different types of complex biological networks.^2^ While the characterization of reference real-world complex networks has uncovered structural similarities among complex networks that are believed to underly their systemic properties,^2, 6^ much less is known about the degree of structural heterogeneity of condition-specific biomolecular networks, and how patterns of variation promote or constrain systems-level behaviors. In cell biology, one intriguing hypothesis is that network heterogeneity emanating from the normal process of development might result in differential behaviors underlying the contrasting cellular phenotypes. In line with this idea, the field of network biology has recently started shifting towards the characterization of condition-specific networks and analysis of circuitry dynamics,^9, 10^ presumably due to the increasing availability of functional genomics and epigenomics assays. For example, Neph and collaborators put forward a methodology to assemble tissue-specific transcription factor networks with the aid of available chromatin accessibility profiles from multicellular genomes.^9, 11–13^ The proposed networks connect each transcription factor (TF) to its incoming TF regulators, thus representing the regulatory structure of the cell in terms of the main regulators (e.g. TFs) and the mutual regulatory interactions among them. More specifically, using digital genomic footprinting (DGF) analysis, TF-TF interactions are established by integrating TF motif matching with DNase I hypersensitive sites (DHS) and high-resolution genomic footprints. Tissue-specificity comes from the condition-specific accessibility of cis-regulatory regions upstream a TF. Using this approach, tissue-specific TF networks have been constructed for model organisms and for human.^9, 14^ Given that the observed TF interactions reflect tissue-specific activity states, we reasoned that the structure and relative systems-level behavior displayed by these networks could provide insights into the biology and differentiation potential of the corresponding tissues.

In order to begin understanding the link between network structure heterogeneity, behavior, and biological phenotypes, here we put forward a computational framework to characterize the structural properties of mammalian tissue-specific TF networks and their behavior, emphasizing the degree of deviation from theoretical expectations. We focus on one systems-level behavior which is informative of the latter: the robustness of the networks to structural perturbations. To this end, we profiled the structural properties of a broad set of TF networks in mouse and human, and we compared the observed behavior across tissues and with expectations from theoretical models. Interestingly, we discovered that embryonic stem cells (ESCs) posses a distinctive regulatory structure: its higher structural similarity to the topological properties expected from a homogeneous network theoretical model endows them with a remarkable resilient behavior. We discuss potential biological implications.

## Results

### Analysis framework

Networks provide a theoretical framework that allows a convenient conceptual representation of interrelations among a large number of elements.^6^ Furthermore, it is usually possible to frame questions about the behavior of the underlying real system by applying well-established measures and analyses over the network representing the empirical data.^15^ Here we focus on tissue-specific networks where nodes represent TFs and links inter-regulatory interactions, and propose an analysis framework with the goal of characterizing the commonalities and differences in behavior against structural perturbations across tissues. We ask whether some tissues display extreme behaviors, and whether or not such deviations an extreme behaviors highlight aspects of the underlying biology. We hypothesize that the differences to be discovered underlie aspects of the observed biological functionality and of the degree of differentiation of the tissues. The proposed framework includes the following steps (see Figure 1). (1) The state of the cell is structurally represented by tissue-specific networks of regulatory interactions among transcription factors as proposed in.^9, 14^ Briefly, a TF is considered regulator of another one when a motif instance of the former TF occurs within a DNase I footprint contained in the proximal regulatory region of the latter TF (10 kb interval centered on the transcriptional start site [TSS]). (2) The system’s behavior of a network is defined as the response of the network against increasing structural perturbations,^2, 16^ and the response is measured by two metrics: the change in giant component size, and the change in efficiency, both relative to the original, unperturbed network (see Methods). The complete behavior is captured by the qualitative properties of the change from start until complete disruption, and we introduce a simple metric to quantify it (Figure 1 a). (3) The structure of each network is numerically characterized by 11 topological measures (Figure 1 b). (4) The degree of deviation of each network relative to expectations from homogeneous (Erdős-Rényi) and heterogeneous (Barábasi-Albert) random graph models is quantified (Figure 1 c).

**Figure 1.**
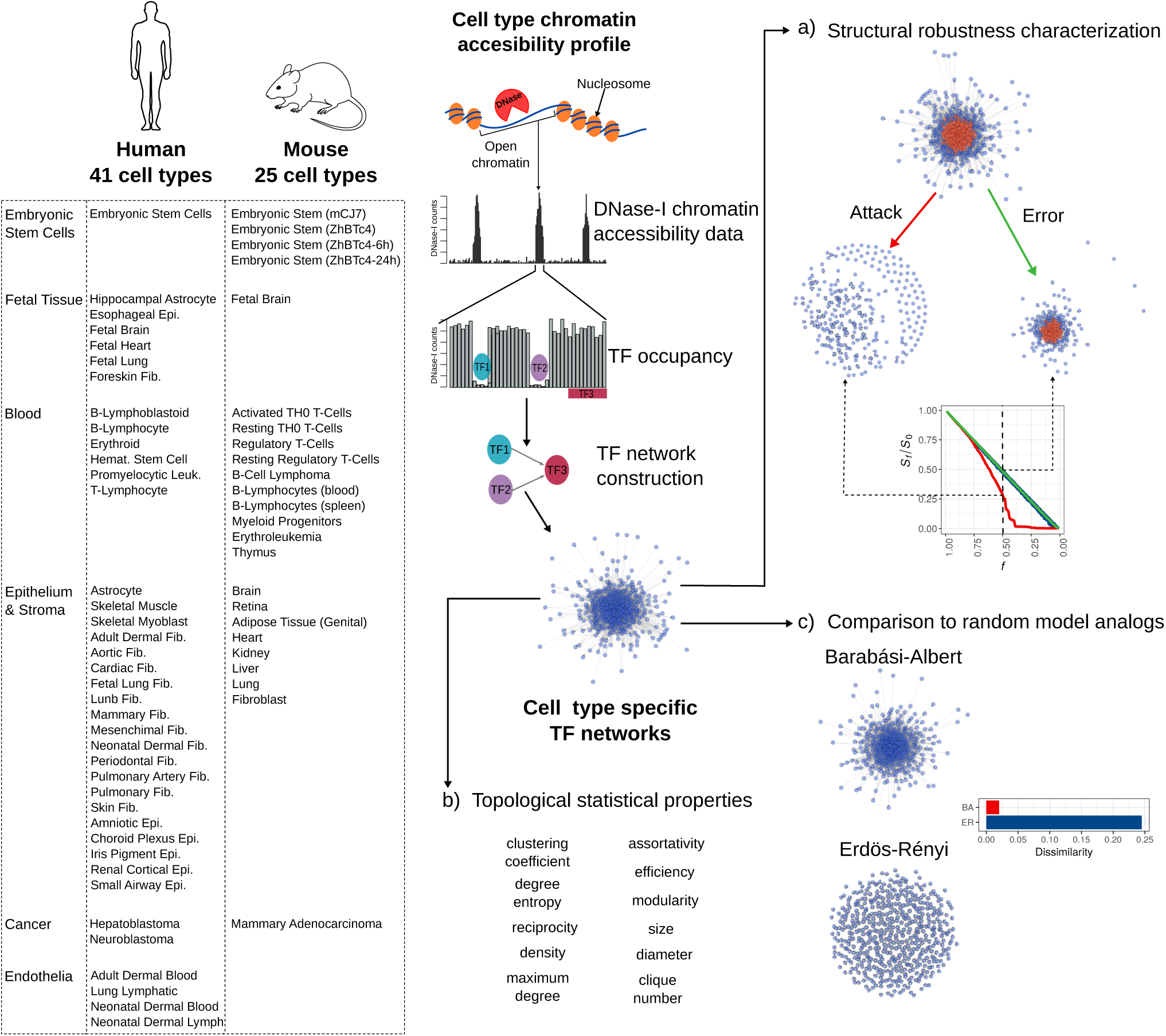
Structural profiling of cell type specific TF networks. a) Structural robustness was measured simulating attacks (removing high degree nodes (red)) and errors (removing randomly selected nodes). b) Networks characterization was done measuring topological features of every network. c) Each network was compared to random model networks by measuring its dissimilarity to an analogous ensemble of homogeneous and scale-free networks.

After applying these steps to each tissue-specific network, we rank the networks based on the robustness of their behavior, we identify those displaying the most extreme response, and we statistically explain the behavior in terms of predictive topological features and relative deviation from analogous homogeneous and heterogeneous random models. Thus, starting from an input set of tissue-specific networks, our framework produces a structural robustness ranking, a set of structural features underlying the behavior, and a mapping of the networks into the homogeneous-heterogeneous network space.

### Network structural differences reveal plasticity of systems behavior upon perturbation

It has been shown that a differential response to random structural perturbations (errors) and directed alterations (attacks) enables a concrete distinction between homogeneous and heterogeneous networks in terms of systems behavior.^16^ A network representing a real complex system is expected to tolerate random failures, but to be more vulnerable against directed attacks targeting key, connected components. Taking this well-established framework, we evaluated the robustness behavior of TF networks across tissues. The operational definition of structural robustness applied here is based on an intuitive idea: disabling a substantial number of nodes will result in an inevitable functional disintegration of a network,^2^ but the degree of tolerance will vary across tissues. We measured tolerance to random perturbations by randomly removing nodes from the networks and quantifying the change in the size of the largest connected component (giant component), and the change in network efficiency – an approximation to loss or gain of network connectivity (see methods). For directed attacks, we repeated the experiments but sequentially removing node in deceasing order of connectivity (degree) (Figure 1 a). We profiled the response to perturbations in 41 human and 25 mouse tissue networks.

Overall, all networks were found to be highly tolerant to random errors. In both mouse and human tissues, the size of the giant component (*S_f_/S*_0_) decreases linearly with *f* without abrupt transitions (Figure 2 a and c, dashed lines). The efficiency of the networks (*E_f_/E*_0_) also shows consistent behavior across all human and mouse tissues: it shows minimal decrease for a large proportion of *f* until it falls abruptly around *f* = 0.8 (Figure 2 b and d, dashed lines). The observed robustness to random failures is consistent with predictions from percolation theory in complex random networks, as it is less likely to perturb key, highly connected components in networks with long-tail degree distribution.^6, 16^ Also consistent with theory, TF networks were found to be much more vulnerable to directed attacks. Interestingly, however, we observed a high degree of variability in the behavior upon attacks across networks. Both measures (giant component size and efficiency) revealed transitions at different fractions *f* of attacked nodes (see Figure 2 a, b, c, and d, solid lines). Interestingly, we found that in both human and mouse the TF networks of embryonic stem cells (ESCs) displayed, relative to other differentiated tissues, an extremely robust behavior against both failure and attack perturbations, the latter being much more pronounced (see Figure 2 a, b, c, and d, red lines).

**Figure 2.**
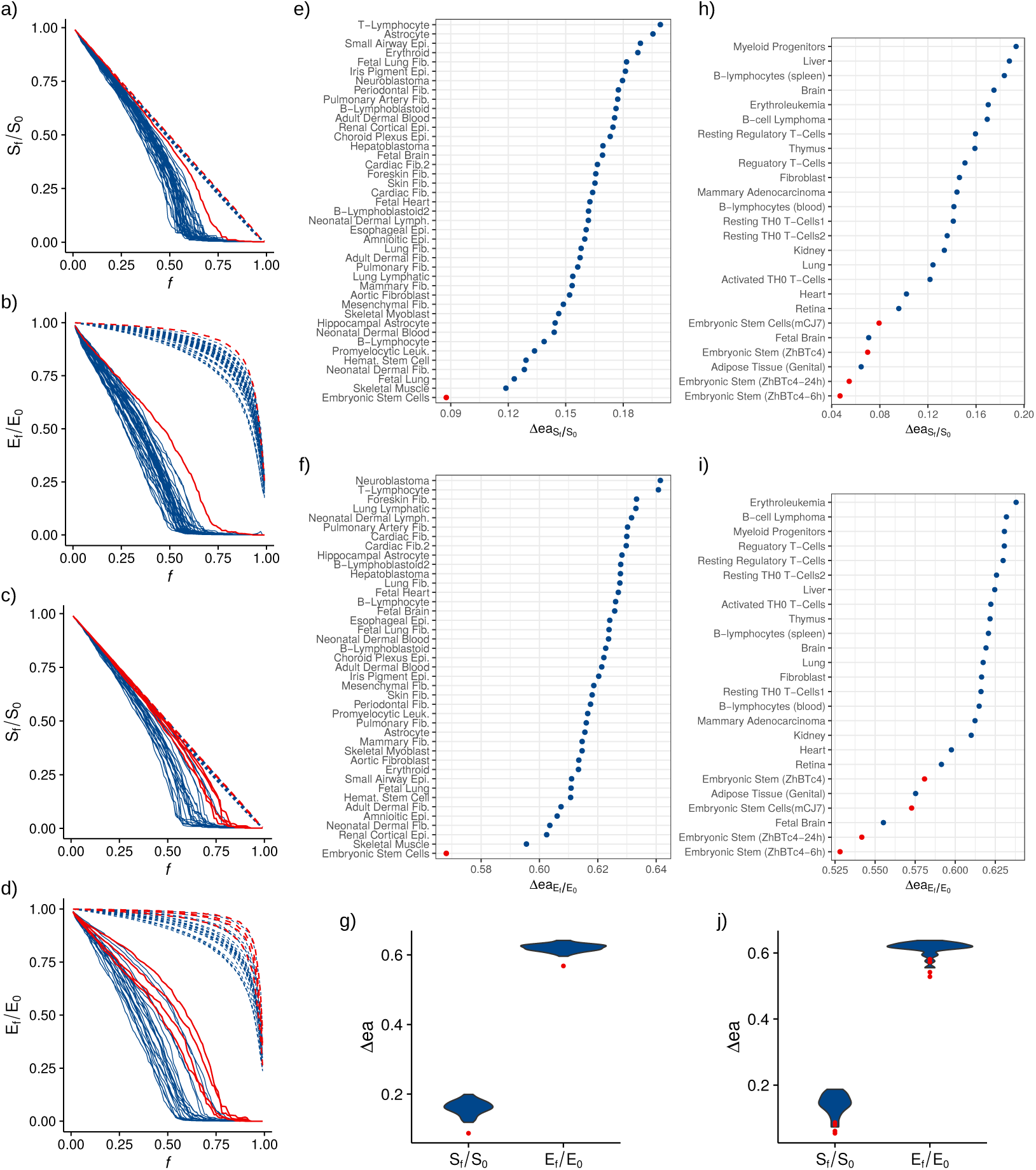
TF networks structural robustness. The behavior against errors (dashed) and attacks (solid) of every cell type is shown, red lines correspond to the ESCs behavior and blue lines to other cell types. a) Human giant component size decrease. b) Human efficiency decrease. c) Mouse giant component size decrease. d) Mouse efficiency decrease. Error-attack deviation measured for each TF network. Human e) giant component and f) efficiency. Mouse h) giant component and i) efficiency. g) Human and j) mouse ∆_*ea*_ distributions, red dots correspond to ESC measurements.

With the goal of quantitatively describing and to analyze the discovered patterns of heterogeneity among tissues, we define the metric error-attack deviation (∆_*ea*_), which simply quantifies the degree of deviation of a given network’s behavior upon directed attack perturbations from that stemming from random errors. We use this metric here as a measure of the structural robustness of complex networks to perturbations, as it reflects the degree to which attacks and errors are tolerated (see Methods). Intuitively, the smaller the value of ∆_*ea*_ the closer the global response of the network against attacks relative to that against error, indicating a higher degree of robustness. We performed the calculation individually for the two damage measures used in this study: *S_f_/S*_0_ and *E_f_/E*_0_. Based on this measure, we can quantitatively compare the differential structural robustness to attacks displayed by the networks. The error-attack deviation of human and mouse cell types corroborates the heterogeneity of structural robustness among cell types, and the extremely deviating behavior of ESCs (Figure 2). ESCs have an error-attack deviation significantly lower than other cell types, highlighting their significantly higher robustness against attacks relative to more differentiated tissues.

### Network structural network rearrangement during differentiation

The observed differences in structural robustness among tissues point to the existence of patterns of variation in global network structure. In order to characterize the structural heterogeneity of TF networks, we analyzed their topology and asked whether specific topological features more predominantly explain the observed robustness patterns. In particular, what structural features underlie the extreme robust behavior of ESCs? As a first approximation we simply asked how similar are the networks to each other? Using a recently published approach to measure network structural dissimilarity (D) (see methods), we computed pair-wise dissimilarity scores for all pairs of TF networks, independently for mouse and human. Network dissimilarity is a useful method for network comparison as it quantifies structural topological differences based on distance probability distributions, capturing nontrivial structural differences^17^ – as opposed to the intuitive counting of presence or absence of common links.

Despite the fact that all TF networks are relatively similar – having average *D* values of 0.040 and 0.064 in human and mouse, respectively – there is variation in the structural similarity among them. *D* ranges from 0.003 to 0.160 in human, and from 0.003 to 0.184 in mouse. Considering pair-wise comparisons in human networks, the ESC is the most dissimilar network for 24 (58.5%) of the tissues. For the remaining 17 tissues, the most dissimilar network corresponds to Astrocyte. These two tissues also have the highest *D* median scores: ESC (0.090) and Astrocyte (0.077). Interestingly, these two networks are also the most dissimilar between one another. Thus, the undifferentiated ESC localizes at one extreme of the topological space while the highly differentiated Astrocyte localizes at the other.

Examining human networks, ESC is clearly different from the other tissues as it is placed in a single branch at the bottom of the distance dendrogram, separated from all the other cell types (Figure 3 a). Mouse networks show a similar pattern to that found in human cell types. Pair-wise comparisons show that the most dissimilar networks are ESCs and the highly differentiated Brain, with these two tissues occupying the extremes in the dissimilarity distribution (Figure 3 b). The mouse ESC ZhBTc-6h has the highest *D* value for 16 of the 25 cell types (64%), while the other two ZhBTc ESCs also rank among the most different networks, and in the remaining 9 cell types the highest *D* value corresponds to Brain. Unbiased hierarchical clustering aggregates the three ESC lines (ZhBTc, ZhBTc-6h, and ZhBTc-24h) in a separate basal branch, together with Genital Adipose Tissue and Fetal Brain. Fetal tissues are expected to display some degree of similarity with ESCs, due to overlap of developmental processes during fetal development. Adipose is an heterogeneous tissue, possibly including undifferentiated adipose stem cells. Overall, the topology of ESCs networks in mouse and human is clearly distinct from adult differentiated tissues such as brain and liver (Figure 3). To further explore the topological differences among tissues, we characterized the structure of every network using 11 standard measures for network topology description (Table 1, see Methods).^6, 7^ These measures capture important characteristics of global structure, which in part determines its functionality. In particular, we seek to dissect the structural heterogeneity among tissues, identify features associated with the observed robustness, and finally map those structural features that discriminate ESCs networks from those of more differentiated tissues.

**Table 1.** Topological features measured for every human and mouse network.

|  | Human Networks |  |  | Mouse Networks |  |  |
| --- | --- | --- | --- | --- | --- | --- |
| Feature | Range | Average | Standard<br>Deviation | Range | Average | Standard<br>Deviation |
| Size | [493, 533] | 521 | 9.27 | [555, 583] | 574 | 7.21 |
| Diameter | [5, 8] | 6.19 | 0.81 | [5, 7] | 5.76 | 0.66 |
| Density | [0.34, 0.68] | 0.051 | 0.007 | [0.47, 0.10] | 0.07 | 0.14 |
| Clique number | [14, 27] | 19.39 | 2.68 | [19, 34] | 26.44 | 3.31 |
| Clustering<br>Coefficient | [0.24, 0.38] | 0.29 | 0.035 | [0.25, 0.36] | 0.30 | 0.026 |
| Assortativity | [-0.21, -0.12] | -0.17 | 0.022 | [-0.18, -0.13] | -0.15 | 0.014 |
| Efficiency | [0.47, 0.54] | 0.51 | 0.016 | [0.50, 0.59] | 0.54 | 0.024 |
| Modularity | [0.10, 0.17] | 0.12 | 0.014 | [0.06, 0.11] | 0.09 | 0.012 |
| Degree<br>Entropy | [5.69, 5.94] | 5.80 | 0.052 | [5.79, 6.09] | 5.93 | 0.078 |
| Reciprocity | [0.03, 0.06] | 0.05 | 0.006 | [0.04, 0.09] | 0.06 | 0.01 |
| Maximum degree | [266, 416] | 348.2 | 35.6 | [334, 574] | 436.9 | 65.2 |

**Figure 3.**
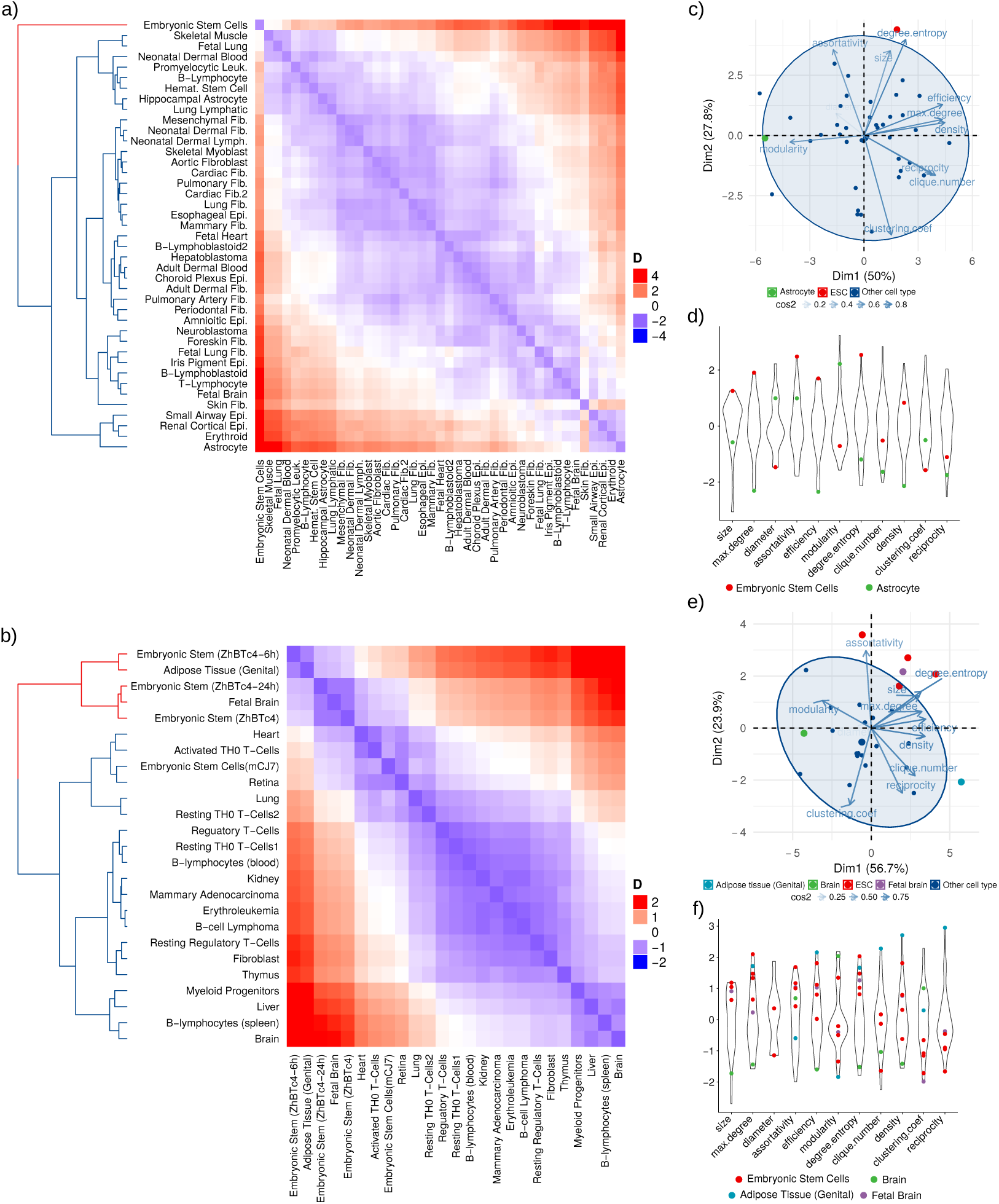
Networks structural profiling. a) and b) Dissimilarity among cell types, heatmaps of scaled *D* values among a) human and b) mouse cell types. c) Human and e) mouse networks topological features PCA. d) Human and f) mouse topological features distribution, colored dots show the value for each feature of the indicated cell type.

In order to explore network aggregation behavior in the feature space, while at the same avoiding collinearity, we performed principal components analysis (PCA), considering the measured topological features. For both human and mouse data, the features with highest contribution for the first principal component (PC) are efficiency, density, maximum degree, and modularity. The features contributing to the second PC are clustering coefficient, assortativity, and degree entropy. Projecting the networks to a 2d space based on PCs, we found no apparent clustering (Figure 3 c and e). However, a closer examination evidences that, as expected, ESCs are separated from the other tissues, having higher values for the second PC. We also highlighted the position of the Astrocyte and Brain networks, considering that these highly specialized tissues are the most structurally different from ESCs. These differentiated networks localize at the other extreme and are characterized for having extremely low values for the first PC. Considering these patterns, ESCs are characterized for having high values of degree entropy, assortativity, and size, but low clustering coefficient. On the other hand, Brain and Astrocyte networks have high modularity, but small density, efficiency, and maximum degree. This pattern is confirmed by the features distribution (Figure 3 d and f).

The topological characterization corroborates the high plasticity of network topologies among tissues, specially an extreme network topology from undifferentiated and highly differentiated tissues. Features analysis shows that tissues spread through features space following two main axes, one going from highly modular to highly efficient networks, and another ranging from highly degree entropic and assortative structures to those with high global clustering. ESCs are among the tissues with more interacting TFs, and these are globally connected in a more promiscuous way, as evidenced by higher levels of entropy in the degree distribution. In contrast, differentiated networks of the Brain and Astrocyte are more structured, as evidenced by low levels of efficiency and density, yet with high modularity. Taking into account the existence of a trade-off between network efficiency and modularity,^18^ this observation hints to a possible path of developmental dynamics of TF network structure in which the system transits from a configuration promoting efficiency in information flow and robustness to a highly modular topology suggestive of functional specialization.

### Interpretation in terms of theoretical network models

As mentioned above, robustness to directed attacks has been linked to homogeneous network topologies, in contrast to the “robust yet fragile” behavior characteristic of heterogeneous (scale-free) networks.^16^ Considering this result, we compared each TF network to analogous ensembles of random homogeneous and scale-free networks generated using the Erdős-Rényi (E-R) and the Barabási-Albert (B-A) models, respectively (see Methods). E-R networks with high number of nodes approach a Poisson degree distribution, symmetric for relatively high average degrees. On the contrary, B-A networks have a characteristic right skewed power-law degree distribution. We compared the real world networks with the theoretical models, with the goal of placing them within a heterogeneity axis by quantifying deviations. Given the discovered high robustness to directed attacks and high degree entropy of ESCs, we reasoned that such a contrast will help clarify the global structural features underlying such behavior.

We measured network structural dissimilarity between each network and its E-R (*D*_*E−R*_) and B-A equivalents (*D*_*B−A*_). As expected, all networks are significantly more similar to B-A than to E-R networks. *D*_*E−R*_ ranges from 0.191 to 0.285 and from 0.139 to 0.287; whereas *D*_*B−A*_ ranges from 0.019 to 0.047 and from 0.013 to 0.055, in human and mouse respectively (Figure 4 d and e). The fact that B-A networks are more similar to the TF networks is consistent with discoveries of other real world complex networks having scale-free topologies.^19^ Interestingly, however, we found clear differences among the networks regarding their relative similarity to each theoretical model. For instance, ESCs have the lowest *D*_*E−R*_ in both human and mouse (Figure 4). In mouse Adipose Tissue and Fetal Brain networks also show low values of *D*_*E−R*_, and ESC line mCJ7 has a *D*_*E−R*_ value higher than the other ESC lines, a pattern already observed in the network topological comparison. In the case of *D*_*B−A*_, a contrasting pattern emerges: ESCs are among the tissues with higher values. Nevertheless, ESC *D*_*B−A*_ values are not significantly different from those of other tissues, falling within the observed distribution of *D*_*B−A*_ (Figure 4 d and e). Considering *D*_*E−R*_ and *D*_*B−A*_ together, ESCs are separated from the other cell types, as shown in Figure 4. A conserved pattern in both human and mouse emerges in which ESCs have a relatively lower dissimilarity to E-R networks and a relatively higher dissimilarity to B-A than the other tissues. Interestingly, mouse ESC line mCJ7, which was separated from the other mouse ESC lines in the cell type dissimilarity histogram (Figure 3 b), clusters together with the other ESC cell lines when taking both *D*_*E−R*_ and *D*_*B−A*_ into consideration.

**Figure 4.**
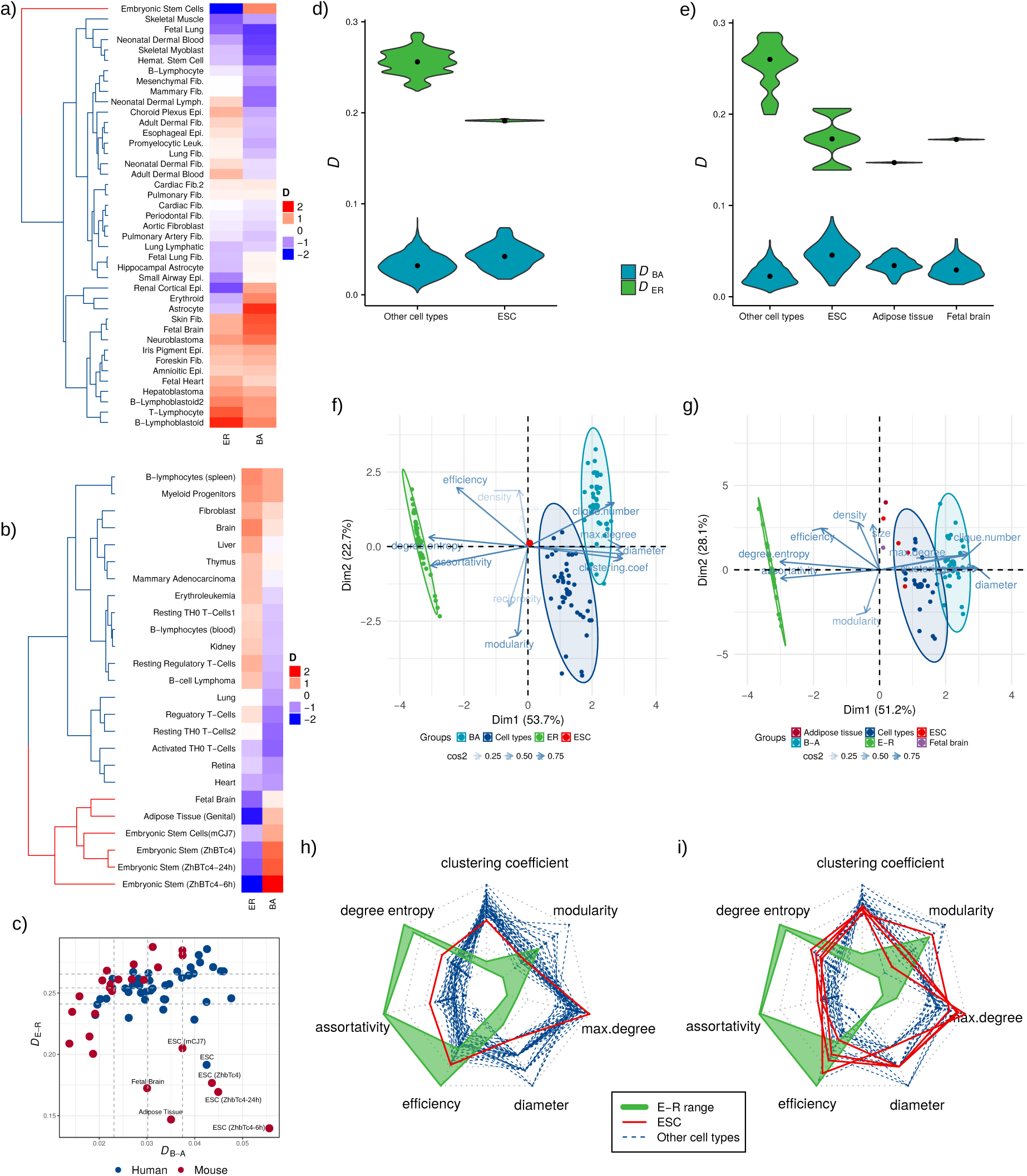
Networks comparison with E-R and B-A model networks. Heatmaps of a) human and b) mouse cell types dissimilarity to model networks. c) Scatterplot of cell types dissimilarity to model networks, dashed lines in both axis correspond to the 25, 50 and 75 percentiles of both measurements. d) and e) Distribution of *D* values for ESCs and other cell types in human and mouse, respectively. Distributions correspond to the dissimilarity with each of the 100 simulated model networks, black dots correspond to distribution median. f) Human and g) mouse network features PCA including real, ER, and BA networks. h) and i) Radar plots of human and mouse networks, respectively, topological features. Green polygons show ER networks’ range for each feature. ESC values are shown in red solid lines, and the other cell types are shown in blue dashed lines.

For every model network we measured the same 11 topological properties we used to characterize cell type networks, and performed a PCA of their features including the real and model networks. In both human and mouse networks, the PCA shows a common pattern. The first component separates three clusters corresponding to each model network and the real networks, situating B-A and E-R networks in the extremes and the real networks between them, closer to the B-A cluster (Figure 4 f and g). As shown in the structural dissimilarity analysis, this pattern confirms that real networks resemble scale-free networks more than homogeneous networks. Real TF networks are situated in between B-A and E-R clusters, thus creating a n space between the two model networks in which real networks can be situated. In both human and mouse, the topological variables with highest contribution to the first PC are degree entropy, assortativity, diameter, and clustering coefficient; together accounting for more than 60% of the variance explained by that component. The pattern shows that E-R networks tend to have higher degree entropy and assortativity, while B-A networks tend to have higher diameter and clustering coefficient (Figure 4 f and g).

We further examined the topological features, looking for those creating an axis between E-R and B-A networks and placing the tissues there. We kept only seven features with the highest contribution to the first two PCs: degree entropy, assortativity, efficiency, clustering coefficient, maximum degree, diameter, and modularity (4 h and i). The feature profile of E-R networks shows high values for efficiency, assortativity, and degree entropy; and relatively low values for diameter, clustering coefficient, and maximum degree. Comparing among tissues, we found that the topological feature profile of ESCs resembles that of E-R networks.

### Network homogeneity predicts structural robustness

The topological analyses we performed show that the topological plasticity of tissue-specific TF networks can be characterized by comparing them to model networks. As mentioned before, this structural differences are associated with the networks’ response to random and directed perturbations. To further understand the structural features underlying the observed structural robustness pattern, we fitted statistical models in an attempt to further uncover explanatory topological features. We defined network structural vulnerability 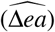 as the mean of the networks’ ∆*ea* for giant component size and efficiency. Using 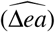 as the response variable, we fitted two statistical models: a linear regression using *D*_*E−R*_ as predictor and a random forest regression using the 11 topological features as predictors. We evaluated each model’s accuracy through five-fold cross validation. The linear regression model has a high predictive power, with cross-validation mean square error of 0.00014. It shows a positive relationship between *D*_*E−R*_ and 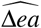 (Figure 5), meaning that the more a network resembles a homogeneous network, the higher its structural robustness. The random forest model also shows high accuracy, with a cross validation mean square error of 0.00025. The most informative topological features are degree entropy and efficiency. Comparing between the models, we see that a simple linear regression has a higher predictive accuracy than the random forest model including 11 topological features. This is reinforced by the fact that the most influential feature of the random forest model is degree entropy, a feature correlated with a network similarity to a homogeneous network, and discovered to characterize ESCs networks. Thus, dissimilarity to E-R model network *D*_*E−R*_, a measure quantifying the degree of homogeneity of a real-world network, is predictive of its structural robustness.

**Figure 5.**
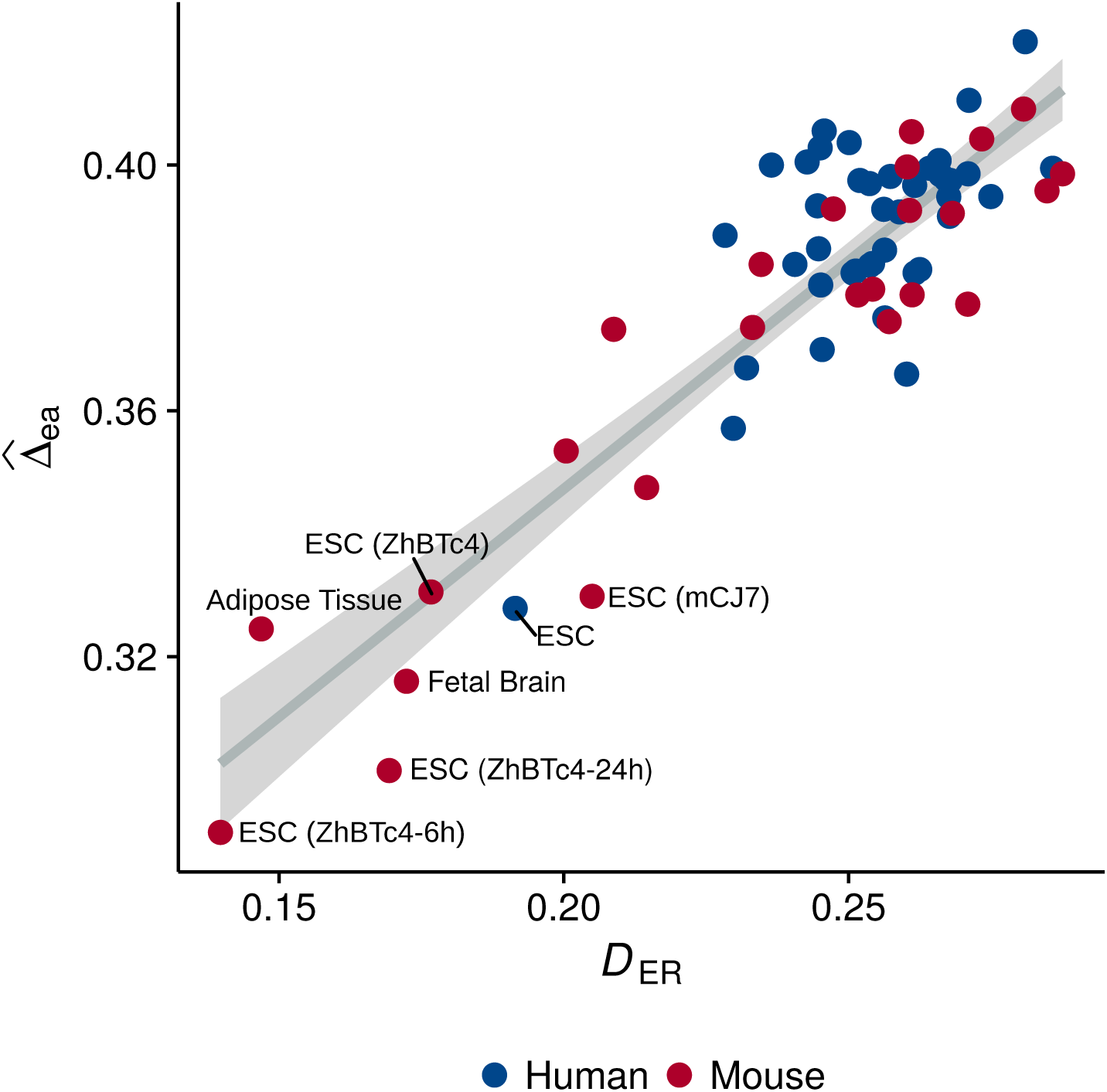
Linear regression. Network vulnerability predicted by networks similarity to Erdős-Rényi model network.

## Discussion

It has been pointed out that insights into the interplay between network structure and dynamics are needed in order to ultimately understand the cell’s functional organization.^2^ Here we studied TF networks’ structure with the goal of better understanding the global behavior of different tissues. As a simple operational approximation, we represented the cell using tissue-specific TF networks. We then analyzed the networks structurally, with the hypothesis that systemic structural differences could provide insights into discriminating systems-level properties ultimately associated with cell behavior. We frame the problem in terms of global structural robustness, a systemic behavior approximated by the vulnerability of networks to both random failure of and directed perturbations.^2, 16^ We found that structural robustness varies significantly across tissues with different levels of differentiation. Interestingly, within the datasets analyzed in both human and mouse, the most robust tissue was also the least differentiated: the embryonic stem cells. Complex network theory has shown the coexistence of extremes in robustness and fragility (”robust yet fragile”) in real-world networks, due to the widespread power-law connectivity distribution associated with complex networks.^16, 20^ The networks underlying ESCs are the most robust against random failure as well as the least fragile against directed attacks, somehow being able to negotiate the observed trade-off between robustness and fragility.

It is known that deviation from the long-tail of theoretical networks with power-law degree distribution reduces the effectiveness of an attack strategy based on targeting the highly connected nodes.^16^ Although all the TF networks analyzed here do have a long-tailed degree distribution, they deviate from theoretical power-law degree distributions (see Supplementary Figures 6 and 7). We analyzed this deviation from a canonical scale-free network by measuring each network’s similarity to theoretical model networks with homogeneous and scale-free topologies. This comparison further exposed the structural heterogeneity among tissues, and the deviating behavior of both undifferentiated (i.e., ESCs) and highly differentiated tissues (i.e., brain tissue and Astrocytes). Furthermore, within the proposed analysis framework, the relative (dis)similarity between a target network and analogous theoretical networks provides insights into the topological characteristics underlying its robustness. For example, the higher structural robustness of ESC networks is explained by its topological resemblance to an Erdős-Rényi homogeneous random network.

**Figure 6.**
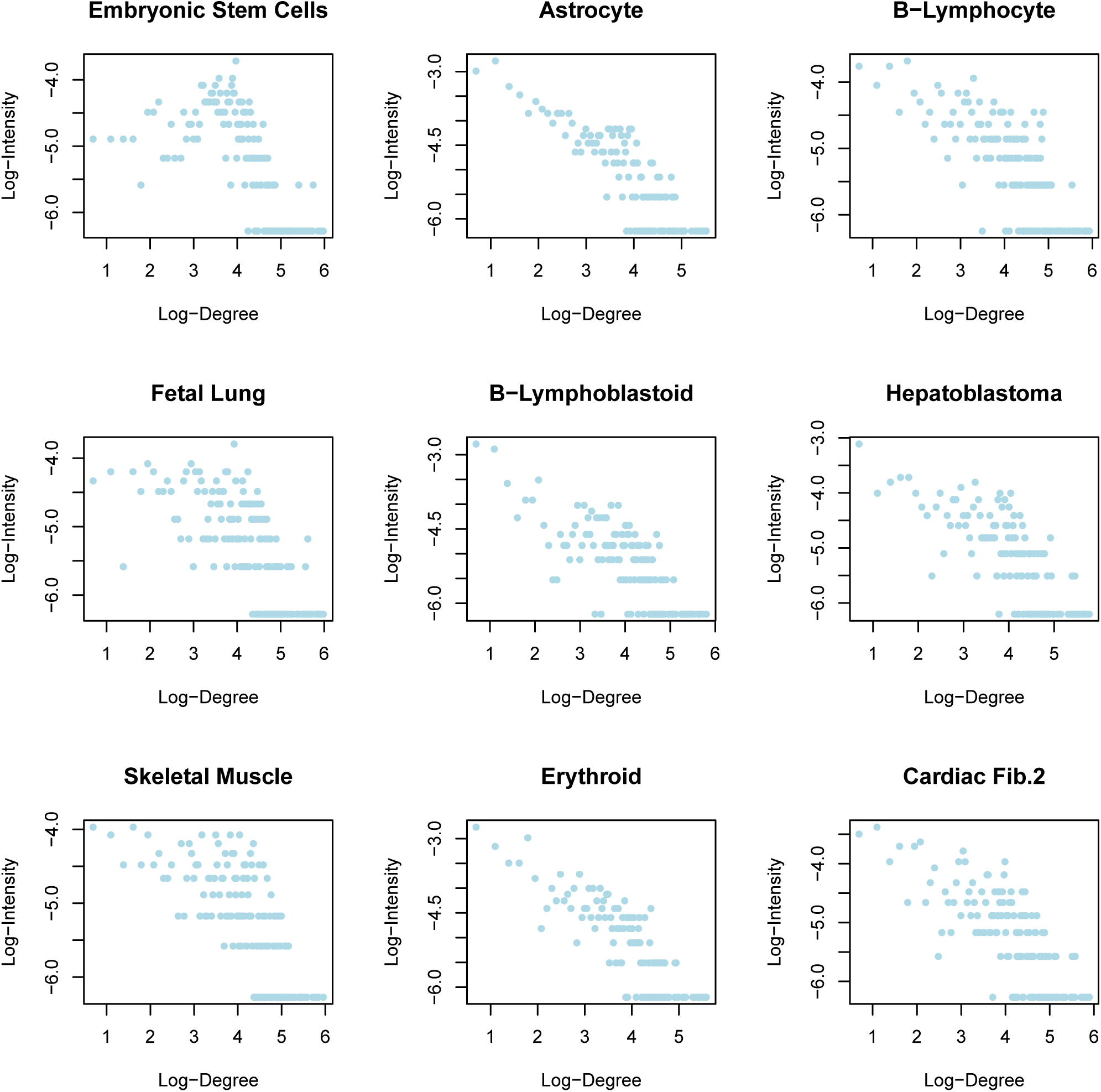
Supplementary 1. Human cell types degree distribution.

**Figure 7.**
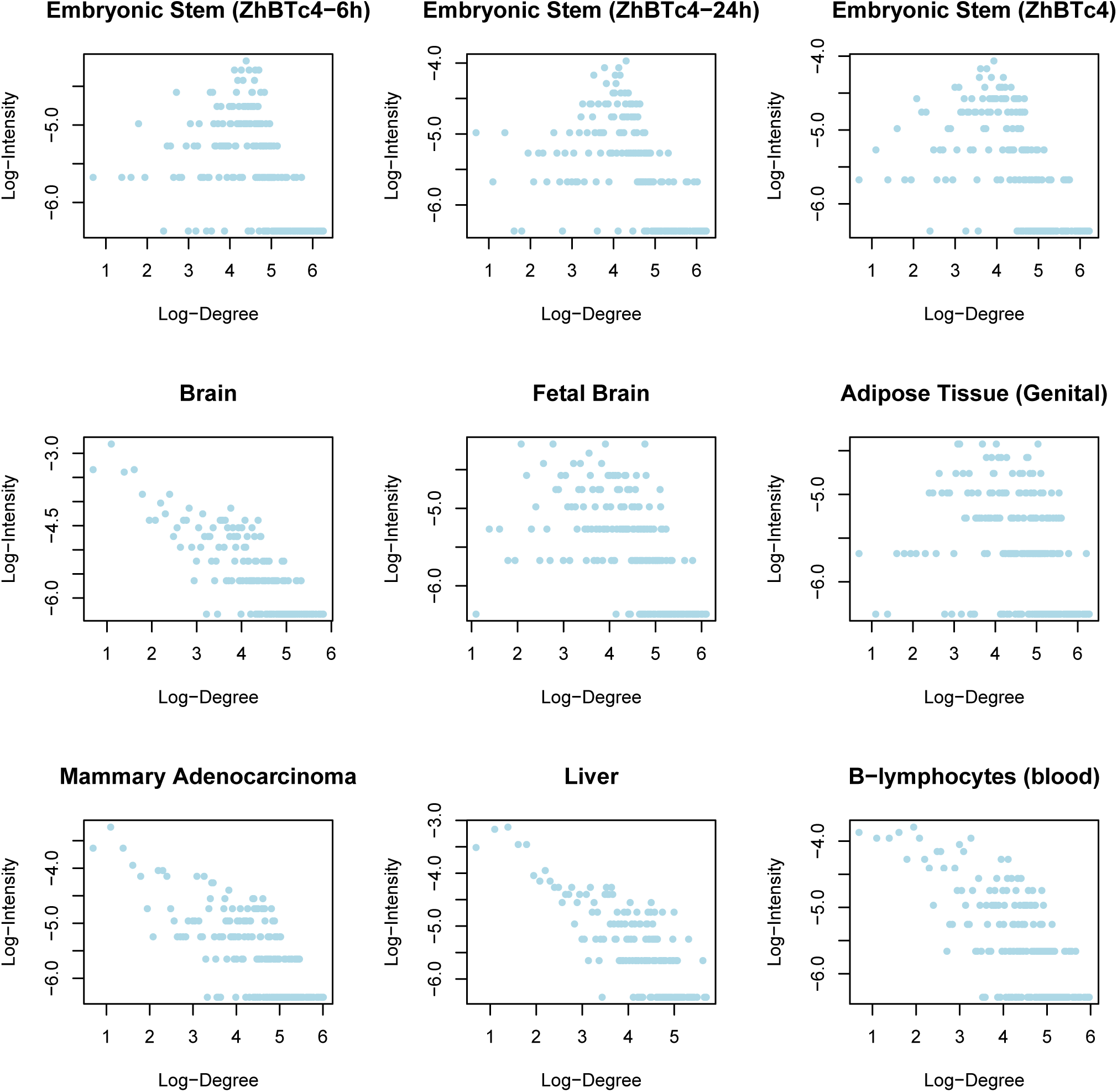
Supplementary 2. Mouse cell types degree distribution.

In terms of biological properties, our results suggest that the ESC state might be able to withstand more and different kinds of errors, due to a more homogeneous network topology. This topological arrangement implies that its main regulator TFs act upon a less constrained chromatin landscape, allowing them to explore it more freely than in differentiated cell types. Consistently, high chromatin accessibility is commonly thought of as a state of loose regulation, and it is associated with stem and dedifferentiated cells – differentiation being considered as a process of increasing chromatin repression.^21^ Another fact adding up to this idea is that ESC have also the highest values of maximum degree among networks, which indicates that the ESC state has more chromatin sites available for a given TF to bind.

In this study we proposed an empirical framework to characterize different networks in terms of their behavior upon structural perturbations and degree of similarity relative to homogeneous and heterogeneous complex networks. It is well know, however, that network topology plays a central role in dynamical behavior. In the cellular context, gene regulatory networks orchestrate cellular behavior.^22^ Theoretical studies have previously analyzed the interplay between structure and dynamics using random Boolean networks.^23^ Networks with a homogeneous topology and relatively high connectivity require fine tuned activation parameters in order to have a stable behavior, and to avoid chaotic dynamics.^23, 24^ This result seems inconsistent with the nature of real biological systems, which have a stable behavior despite fluctuations in surrounding environmental parameters. In other words, resilience is a characteristic of biological systems. Interestingly, for networks with a scale-free topology stable behavior emerges without the fine tunning requirement.^23, 24^ Considering our results in this structure/dynamics context, similarity to homogeneous networks found in the ESC networks analyzed here is likely to produce less ordered dynamics than more differentiated tissues, which, at the same time would allow them to explore more freely the state space and to reach multiple different network states. Interestingly, this view is consistent with the observed high heterogeneity in gene expression and the balance between robustness and plasticity characteristic of ESCs.^25–28^ Although we did not consider dynamical analysis in this study, but rather limited ourselves to the structural characterization of the networks and their behavior, disentangling structure and dynamics will be the focus of future work.

Summarizing, in light of the amount of data on biological interactions being generated in the post-genomic era, a systems level perspective is required to gain understanding of the biological systems as a whole. Our structural analysis of tissue specific TF networks aims at that objective, trying to find the connections between transcriptional networks structural heterogeneity and biological phenotypes. Our treatment of structural robustness as a network systems-level behavior revealed differences among cell types that could be furthered dissected through topological analyses. We want to stress the applicability of our comparison of real world complex networks not only for a structural characterization, but also as an approximation to their possible dynamic behaviors. Finally, the empirical analysis framework proposed here can be applied to any set of related networks whose structural heterogeneity is suspected to underly differential real life behavior.

## Methods

### Transcription Factor Networks

Human and mouse transcription factor networks (TFNs) were constructed based on DNase-seq data and digital genomic footprinting as shown in.^9, 14^ Human networks set include 41 distinct cell and tissue specific networks composed of 493 to 533 sequence-specific transcription factors. Mouse networks set include 25 cell and tissue specific networks composed of 555 to 583 sequence-specific transcription factors. For simplicity, we use the term tissue-specific through the text to refer to both cell type and tissue. Network data were downloaded from http://www.regulatorynetworks.org/. Most current versions for human (v09162013) and mouse (v12032013) were used.

### Modeling topological robustness

Topological robustness was approximated by profiling the network’s behavior in response to random and directed structural perturbations. Site percolation was used as a process to model component failure using computer simulations.^6^ Increasing fractions of a network’s vertices were removed, along with the edges connected to those vertices. Following^6, 29^ a percolation process was considered in the general sense – i.e., including different ways of vertex removal. The error experiments performed correspond to the simplest percolation process where a fraction of vertices was chosen uniformly at random and removed. For every network, error experiments were repeated 1000 times and the mean error behavior was calculated. Directed (Attack) experiments were simulated by removing vertices in decreasing order of centrality based on vertex degree. Nodes were progressively removed from one to a hundred percent of nodes.

### Quantifying network structural robustness

Two quantitative measures of network damage were used to characterize the phenomenology associated to the damage process applied to each TF network. As a first approximation, the macroscopic (systemic) behavior of the networks in response to damage was characterized by the evolution of the giant component size relative to its initial value as a function of the fraction of removed vertices *f* (*S_f_/S*_0_). As an additional approximation, the global efficiency *E* of a network was used to quantify how communication becomes less efficient as damage increases, this measure was also calculated relative to its initial value and as a function of the fraction of removed vertices *f*. The latter measure assumes that the efficiency for sending information between two vertices *i* and *j* is proportional to the reciprocal of their distance, and is calculated as follows:^7, 8^

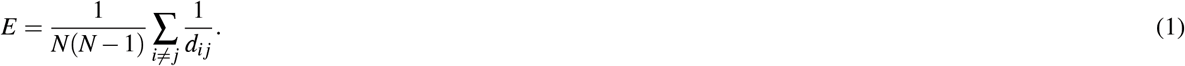

The measure *E* corresponds to the average inverse geodesic length – i.e., the harmonic mean of the geodesic distances:^7^

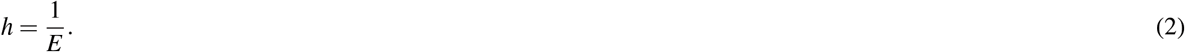

### Error-Attack Deviation

The measure error-attack deviation ∆_*ea*_ introduced herein, was used to quantify the degree of robustness to attacks relative to that against errors. The metric is simply the root mean square deviation between the observed error and the attack behaviors:

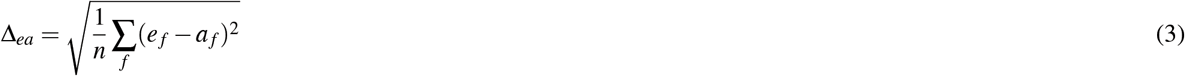

where *e*_*f*_ (*a*_*f*_) represents the a normalized measured of damage behavior under the random or (directed) removal of a fraction *f* of nodes. In this study *S_f_/S*_0_ and *E_f_/E*_0_ were used as damage measures (see Results).

### Networks Topological Characterization

Networks’ topology was analyzed by quantifying topological dissimilarity and measuring 11 structural features commonly used in complex network theory.^6, 7^

#### Network dissimilarity

Network dissimilarity measurement was done following the approach proposed by Shieber et al.^17^ This method compares networks topology based on quantifying differences among node distance probability distributions, representing all nodes connectivity distances, extracted from the networks. It returns non-zero values only for non-isomorphic graphs, and quantifies structural topological differences that have an impact on information flow through the network. We measured network dissimilarity following the algorithm proposed in,^17^ using the suggested parameters.

#### Networks structural characterization

We described networks’ topology by measuring 11 features: number of nodes, diameter, maximum degree, density, clustering coefficient, assortativity, efficiency, modularity, degree entropy, clique number, and reciprocity. Following the measurement definitions in.^7^

#### Null models

To compare cell type networks with random models, we generated random networks with the same number of nodes and links. Two sets of random networks were created: one set following Erdő s-Rényi model (E-R networks) with exponential degree distribution, and the second set following Barabási-Albert model of growing networks with power-law degree distribution (B-A networks). In order for the B-A networks to have an equivalent number of edges to its real counterpart, the number of outgoing edges added to each new node in the network was taken from the out degree distribution of the real network.

For each real network, 100 E-R and B-A random networks were created. Every random network was structurally characterized measuring the 11 topological features measured in the real networks, and dissimilarity to its real equivalent was quantified. Mean values for the dissimilarity and topological features were estimated for each ensemble of random networks.

#### Predictive modeling

Predictive models were fitted using networks’ vulnerability as a response variable and structural features as predictors. We defined network vulnerability 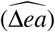 as the mean between error-attack deviation to giant component size and efficiency:

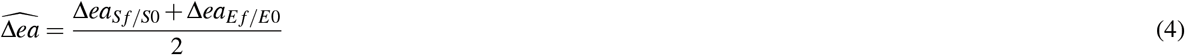

The first model we fitted was a linear regression predicting 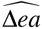 from the network’s mean dissimilarity to its E-R analogs (*D*_*E−R*_). The second model we fitted was a random forest regression, predicting 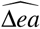 from the 11 topological features measured above, this model was was created with 1000 trees. Features’ influence on the random forest model was measured by the mean decrease in mean square error. As a way to evaluate the models’ accuracy, we performed a five-fold cross validation of both models, keeping the test mean square error as accuracy measurement.

### Implementation

All the methods presented here were implemented using the *R* statistical programming environment (www.R-project.org) and the igraph package.^30^

## Acknowledgements

This work was supported by Consejo Nacional de Ciencia y Tecnología, (CONACYT: 240180, 180380, 2015-01-687) and UNAM-DGAPA-PAPIIT (IN211516, ININ208517, IN205517, IN204217). J.C.P. is a doctoral student from Programa de Doctorado en Ciencias Biomédicas, Universidad Nacional Autónoma de México (UNAM) and received fellowship 446988 from CONACYT. We thank Diana Romo for logistic support.

## Author contributions statement

J.D.V designed and coordinated the study. J.C.P conducted the analyses. J.C.P and J.D.V analyzed the results, and wrote the manuscript. E.A.B provided resources and discussed the problem and results. All authors read and approved the manuscript.

